# Plasticity in female timing may explain current shifts in breeding phenology of a North American songbird

**DOI:** 10.1101/2021.09.29.462407

**Authors:** Abigail A. Kimmitt, Daniel J. Becker, Sara N. Diller, Nicole M. Gerlach, Kimberly A. Rosvall, Ellen D. Ketterson

**Affiliations:** Department of Biology, Indiana University, 1001 E. Third St., Bloomington, Indiana 47405; Department of Ecology and Evolutionary Biology, University of Michigan, 1105 North University Ave, Ann Arbor, MI 48109; Environmental Resilience Institute, Indiana University, 717 E. Eighth St., Bloomington, Indiana 47408; Department of Biology, University of Oklahoma, 730 Van Vleet Oval, Norman, OK 73019; Department of Biological Sciences, Western Michigan University, Kalamazoo, MI 49008; Department of Biology, University of Florida, P.O. Box 118525, Gainesville, FL 32611

**Keywords:** climate change, timing of breeding, selection, phenotypic plasticity, phenological shifts, bird

## Abstract

1. Climate change has driven changes in breeding phenology. Identifying the magnitude of phenological shifts and whether selection in response to climate change drives these shifts is key for determining species’ reproductive success and persistence in a changing world.
2. We investigated reproductive timing in a primarily sedentary population of the dark-eyed junco (*Junco hyemalis*) over 32 years. We predicted that juncos would breed earlier in warmer springs in response to selection favouring earlier breeding.
3. To test this prediction, we compared the annual median date for reproductive onset (i.e., egg one date) to monthly spring temperatures and examined evidence for selection favouring earlier breeding and for plasticity in timing.
4. Egg one dates occurred earlier over time, with the timing of breeding advancing up to 24 days over the 32-year period. Breeding timing also strongly covaried with maximum April temperature. We found significant overall selection favouring earlier breeding (i.e., higher relative fitness with earlier egg one dates) that became stronger over time, but strength of selection was not predicted by temperature. Lastly, individual females exhibited plastic responses to temperature across years.
5. Our findings provide further evidence that phenotypic plasticity plays a crucial role in driving phenological shifts in response to climate change. For multi-brooded bird populations, a warming climate might extend the breeding season and provide more opportunities to re-nest rather than drive earlier breeding in response to potential phenological mismatches. However, as plasticity will likely be insufficient for long-term survival in the face of climate change, further research in understanding the mechanisms of female reproductive timing will be essential for forecasting the effects of climate change on population persistence.

## 1. Introduction

Climate change is greatly affecting plant and animal life (Root et al. 2003; Scheffers et al. 2016; Staudinger et al. 2013). Phenological shifts are common (Piao et al. 2019; Scheffers et al. 2016), suggesting that many species can adjust to climate change (Charmantier and Gienapp 2014; Saalfeld and Lanctot 2017). Identifying the magnitude of phenological shifts and their selective drivers in response to climate change are key for conservation efforts (Charmantier & Gienapp 2014). Numerous studies have investigated phenological shifts in passerine birds in the last two decades; however, the number of long-term datasets for unique species that can account for breeding timing as well as reproductive success is limited (i.e., <10 passerine species). Additionally, these species greatly vary in life history (e.g., migratory strategy [migrant vs. resident], breeding duration [single-vs. multi-brooded], nesting strategy [cavity vs. open cup nesting], diet, and habitat) and geography, all of which could directly affect selection pressures on breeding phenology (Dunn & Møller 2014). Investigating the potential drivers of phenological shifts in additional species with distinctive life histories will allow for more accurate predictions of which populations will be able to adapt to the changing climate. To date, many studies investigating phenological shifts in birds have been focused on European species, with some work in North America species that are predominantly migratory, but see (Wilson *et al*. 2007; Watts *et al*. 2019). In this study, we contribute to this growing body of literature by analysing a long-term data set of the breeding efforts of a North American, resident songbird population.

The relative role of microevolutionary change versus behavioural plasticity in driving phenological shifts remains under debate (Charmantier & Gienapp 2014), as it likely varies across species in relation to their life history and their ability to adapt to climate change. Climate change could drive directional selection favouring earlier breeding in birds by influencing phenology in related trophic levels (e.g., prey, competitors), thus affecting offspring survival (Charmantier & Gienapp 2014). Evolutionary adaptation can then occur if directional selection favouring earlier breeding acts on heritable traits with genetic variation (Hoffmann & Sgrò 2011). Phenological changes may also reflect behavioural plasticity, or the ability of an individual to modify its behaviour based on the environment (Sih *et al*. 2010; Van Buskirk, Mulvihill & Leberman 2012; Beever *et al*. 2017). Behavioural plasticity and its underlying mechanisms can allow individuals to respond more quickly to a changing climate, as compared to microevolutionary change in response to selection (Sih *et al*. 2010; Charmantier & Gienapp 2014; Beever *et al*. 2017). However, plasticity is not always adaptive (Duputié *et al*. 2015) and is unlikely to be sufficient to allow populations to respond long-term to climate change (Ghalambor *et al*. 2007; Gienapp *et al*. 2013).

Here, we used long-term data collected from Dark-eyed Juncos (*Junco hyemalis*), a north-temperate sparrow found in Canada and the United States, to investigate changes in their breeding phenology. Juncos serve as a model songbird species for studies of ecology and evolution (Ketterson & Atwell 2016). Specifically, we focused on a breeding population of Carolina Dark-eyed Juncos (*J. h. carolinensis*) that resides in the Appalachian Mountains year-round, with some individuals migrating short distances (e.g., altitudinal migrants). We first asked whether median monthly air temperature in early spring changed over the 32-year study, predicting that spring temperatures would increase over time. We next compared annual average egg one dates (i.e., initiation of breeding, or the date of first egg laid in the year) to spring temperatures over time and predicted that egg one dates would be earlier over time in response to a warming climate. We also asked whether selection acted on earlier breeding by assessing the relationship between female annual relative fitness and egg one date. We then used a model comparison framework to identify climatic drivers of the strength of selection across our study period. We predicted that selection would favour earlier breeding, especially in warmer springs. Lastly, we used a random regression model approach to evaluate the degree of female plasticity in response to spring temperatures. We predicted that individuals that bred in multiple years would vary their initiation of breeding in response to changing spring temperatures proportional to annual differences in temperature.

## 2. Methods

### **a)** Study system and breeding data

Since 1983, a breeding population of Dark-eyed Juncos has been monitored at Mountain Lake Biological Station (MLBS) and the surrounding Jefferson National Forest (37°22’N, 80°32’W) (Chandler *et al*. 1994). At the beginning of each breeding season (April-May), birds on the study site were caught using mist nets and Potter traps and banded with a unique USFWS metal band and distinctive combinations of colour bands. Researchers searched for nests every year, identifying parents and tracking the progress of the nest. Egg one date, expressed as Julian date, was observed directly, or for nests found after the start of egg-laying, was calculated based on the day nestlings hatched or left the nest (Nolan *et al*. 2002). Breeding data from 1983–2015 were used for this study except for 2013 due to limited research effort. Records where female ID or egg one date were unknown were removed. Female subjects that were implanted with exogenous testosterone during a five-year study were (Clotfelter *et al*. 2004; Ketterson, Nolan & Sandell 2005) were also removed.

To calculate true egg one dates, we excluded any known re-nests. Also, knowing that the first nest found for a female might not be her true first nest, we eliminated nests whose egg one dates came later than each year’s median egg one date from known re-nests. Our data filtering resulted in 1,244 first nests of 936 female juncos between 1983 and 2015. Annual differences in research effort (number of nests found) did not explain variation in egg one dates (see Supplementary Materials; Fig. S1).

Because the distributions of egg one dates were not normal in some years, we calculated median annual egg one dates from first nests. Using both first nests and renests for each year, we calculated the annual total number of eggs and total number of fledglings produced by each female. Females were grouped into two age classes based on plumage (Pyle 1997) or records from previous breeding seasons: second years (SY; first breeding season) and after second years (ASY; second or later breeding season). Finally, since most open-cup nests fail due to predation (Ricklefs 1969), we estimated annual predation rates of nests by calculating the annual percentage of nests that failed at the egg or nestling stage before fledging.

### b) Temperature data

Between November 16, 1971 and January 31, 1998, temperature data (daily minimum; T_in_ and maximum temperature; T_max_) were collected from MLBS via a National Oceanic and Atmospheric Administration (NOAA) weather station (Network ID GHCND: USC00445828, hereafter, “Logger A”). On June 24, 1994, a second data logger (Campbell CR10) was established at MLBS that records temperature every half hour. To permit comparing data across devices, we calculated daily T_max_ and T_min_ from this MLBS data logger (hereafter, “Logger B”). From T_min_ and T_max_, we calculated daily median temperature (T_med_) for both loggers. Since the two weather stations overlapped from 1994-1997, we confirmed that Datasets A and B were strongly correlated and then combined the datasets (*see Supplementary Materials)*.

Monthly average T_min_, T_max_, and T_med_ were calculated for March–May for each year. Data were available for all years (1983-2015), except for missing March and April data for 1991 and 2002 and missing March data for 2004.

### c) Temporal patterns

All statistical analyses were conducted in R (version 4.0.0). Temperature is not expected to exclusively change linearly over time, so we first fit generalized additive models (GAMs) with a smooth term for year to flexibly determine temporal trends in average T_min_, T_max_, and T_med_ during spring (March-May) when birds were initiating breeding.

We fit a GAM with a smooth term for year to first investigate change over time in median egg one date. Most female juncos lay their first egg in late April-early May, and the final stages of reproductive development can take anywhere from days to weeks (Williams 2012), such that temperatures prior to laying likely have the greatest influence on female reproductive timing. Therefore, to determine how annual spring temperatures and temporal variation related to median annual egg one date, we fit independent GAMs with smooth terms for both year and each of 9 temperature variables (average T_med_, T_min_, and T_max_ for March, April, and May). We then compared model fit using Akaike information criterion corrected for small sample size (AICc) and Akaike weight (*w*_*i*_; Burham & Anderson 2002). We considered models within two ΔAICc of the top model to be competitive. All GAMs were fit with the *mgcv* package using a Gaussian distribution and thin plate splines (Wood 2017). We used maximum likelihood (ML) for model selection but refit all final GAMs using restricted ML (REML). We used the most competitive temperature covariates in our subsequent selection and plasticity analyses.

### d) Selection analyses

Selection acting on start of breeding was defined as the slope of a regression of relative fitness (i.e., total number of fledglings per year per female divided by annual population mean total fledglings) on egg one date (Lande & Arnold 1983). We adjusted for age, annual total eggs per female, and predation rate when estimating selection acting on egg one date by including these as fixed effects (Marrot, Garant & Charmantier 2017). As relative fitness was zero inflated, we used compound Poisson generalized linear models (GLMs) or generalized linear mixed models (GLMMs) with the *cplm* package (Zhang 2013); however, we also estimated the linear selection gradient using a conventional LMM (Lande & Arnold 1983). We ran analyses on egg one dates standardized annually (zero mean and unit variance) to control for environmental covariance between fitness and this trait across years (Kingsolver *et al*. 2001; Marrot *et al*. 2018). Owing to missing values, these fitness analyses included 1,182 first nests from 898 females. We included year and female ID as random intercepts to control for multiple observations per year and females that bred more than one year.

We ran selection analyses and model comparisons using two approaches. First, we derived selection gradients on a per-year basis using GLMs. We then compared univariate linear regressions with each of our primary temperature variables identified from our GAM analyses using AICc and Akaike weights (Burham & Anderson 2002). To account for uncertainty in estimates of selection gradients, these models included weighting by the annual sample size, (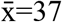 first nests ± 2 SE). Next, we tested for relationships between temperature and the strength of selection on egg one date at the individual level using GLMMs fit to our full dataset. We compared variants of our base selection GLMM with an interaction between egg one date and each temperature variable, again using AICc and Akaike weights. We derived a marginal and conditional *R*^*2*^ as measures of fit (*R*^*2*^_*m*_ and *R*^*2*^_*c*_; (Nakagawa, Johnson & Schielzeth 2017). Owing to missing temperature data, we used a reduced dataset of 1,123 first nets from 871 females to compare GLMMs.

### e) Behavioural plasticity

To assess the degree of individual female plasticity in egg one date in relation to spring temperature, we used a random regression model (RRM) approach (Nussey, Wilson & Brommer 2007). RRMs are a particular case of GLMMs where individuals vary in the elevation (i.e., intercept) and slope of their reaction norms. For females that bred in at least three years of our study (*n*=62 individuals representing 206 egg one date observations, hereafter “returning females”), we fit a RRM with a fixed effect of temperature, a random intercept of female ID, and their interaction (i.e., a random slope) using ML with the *lme4* package (Pinheiro & Bates 2006). We again included age, annual total eggs per female, and predation rate as covariates and included an additional random intercept for year. We again compared among temperature predictors using AICc and Akaike weights and refit RMMs with REML to derive *R*^*2*^_*m*_ and *R*^*2*^_c_ and estimate variance for random effects. We then used sequential likelihood ratio tests to assess if random intercepts and slopes were significantly different from zero (i.e., denoting significant inter-individual variation in reaction norms) (Nussey *et al*. 2005). To visualize individual female slopes, we held each female at the ASY age class at its mean annual clutch size and annual predation rate.

## 3. Results

### **a)** Spring temperatures

March average T_med_, T_min_, and T_max_ did not significantly change between 1983 and 2015 at MLBS. April average T_med_ and T_min_, but not T_max_, significantly increased over time. Finally, average May T_min_ significantly increased over time, but T_med_ and T_max_ did not (Table S1; Fig. 1).

**Figure 1.**
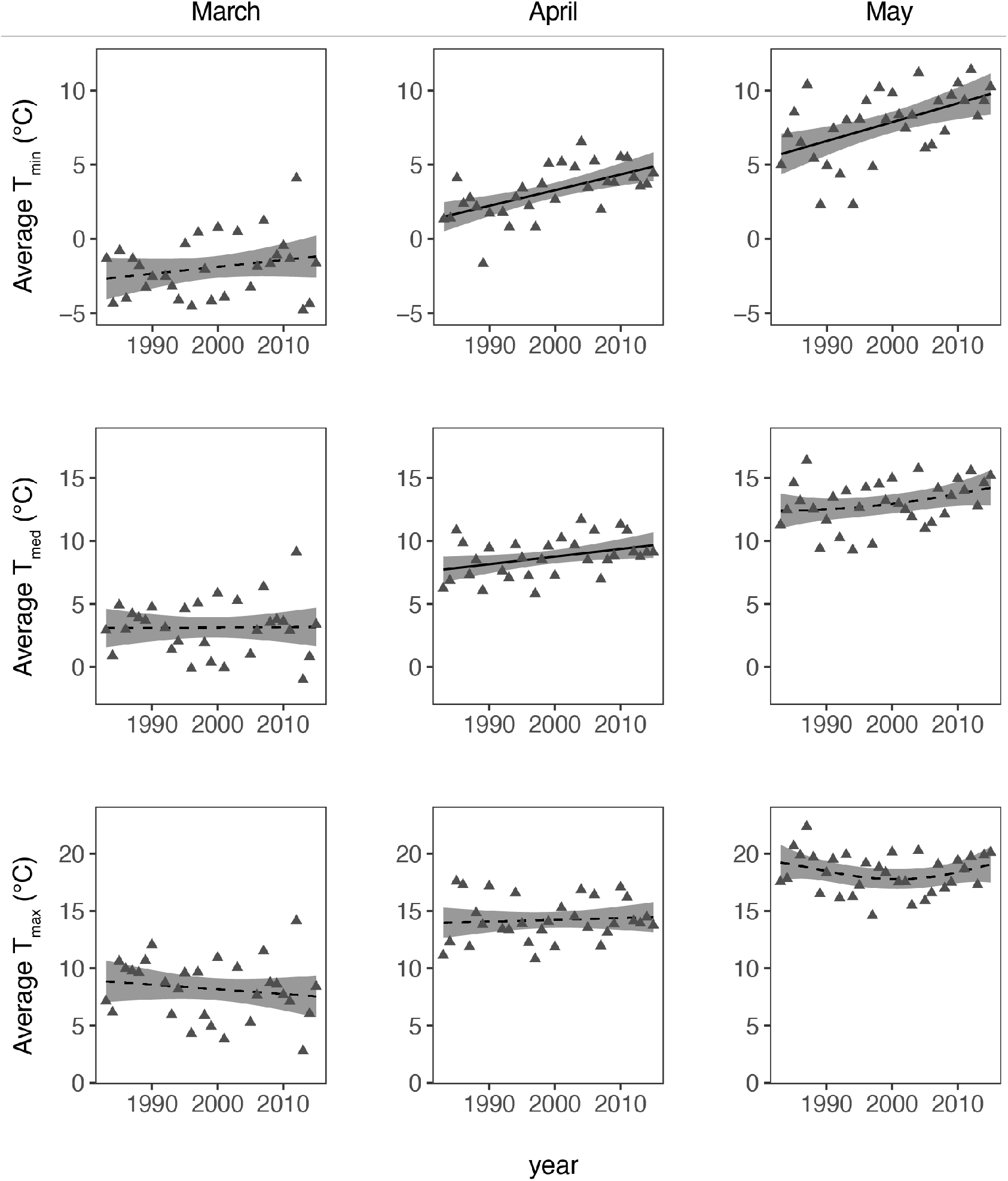
Independent relationships between year and median, minimum, and maximum temperatures in March-May from 1983 to 2015. All prediction lines and confidence bands from the GAMs are created as a function of year as a smooth term and overlaid with original data. Average April T_med_, April T_min_, and May T_min_ change significantly over time (April T_med_: *F*_*1,1*_= 4.79, *p*= 0.037, *R*^*2*^= 0.11; **April T**_**min**_: *F*_*1,1*_= 15.86, *p*< 0.001, *R*^*2*^= 0.33; **May T**_**min**_: *F*_*1,1*_= 11.41, *p*= 0.002, *R*^*2*^= 0.25)

### b) Timing of reproductive onset

Median egg one date varied significantly over time (*F*_*1,1*_*=* 4.43, *p=* 0.044, *R*^*2*^= 0.10), with a 13-day difference from the first year (May 19, 1983) to the final year (May 6, 2015; Fig. 2A). Considering the largest difference in breeding phenology over the 32 years, females advanced egg one dates up to 24 days over the study (range=April 30-May 24). When testing effects of monthly temperatures after accounting for nonlinear effects of year, the best GAM included April average T_max_ (*w*_*i*_*=* 0.97; Table S2). In this model (*R*^*2*^ =0.65), egg one date was predicted by the average April T_max_ (*F*_*1*.*7, 2*.*1*_ *=* 14.84, *p<* 0.0001) and year (*F*_*2*.*4, 3*.*0*_*=* 5.28, *p=* 0.007) (Fig. 2B). Since an April temperature was the best predictor of lay date, we proceeded using only the three April temperature variables for selection and plasticity analyses.

**Figure 2.**
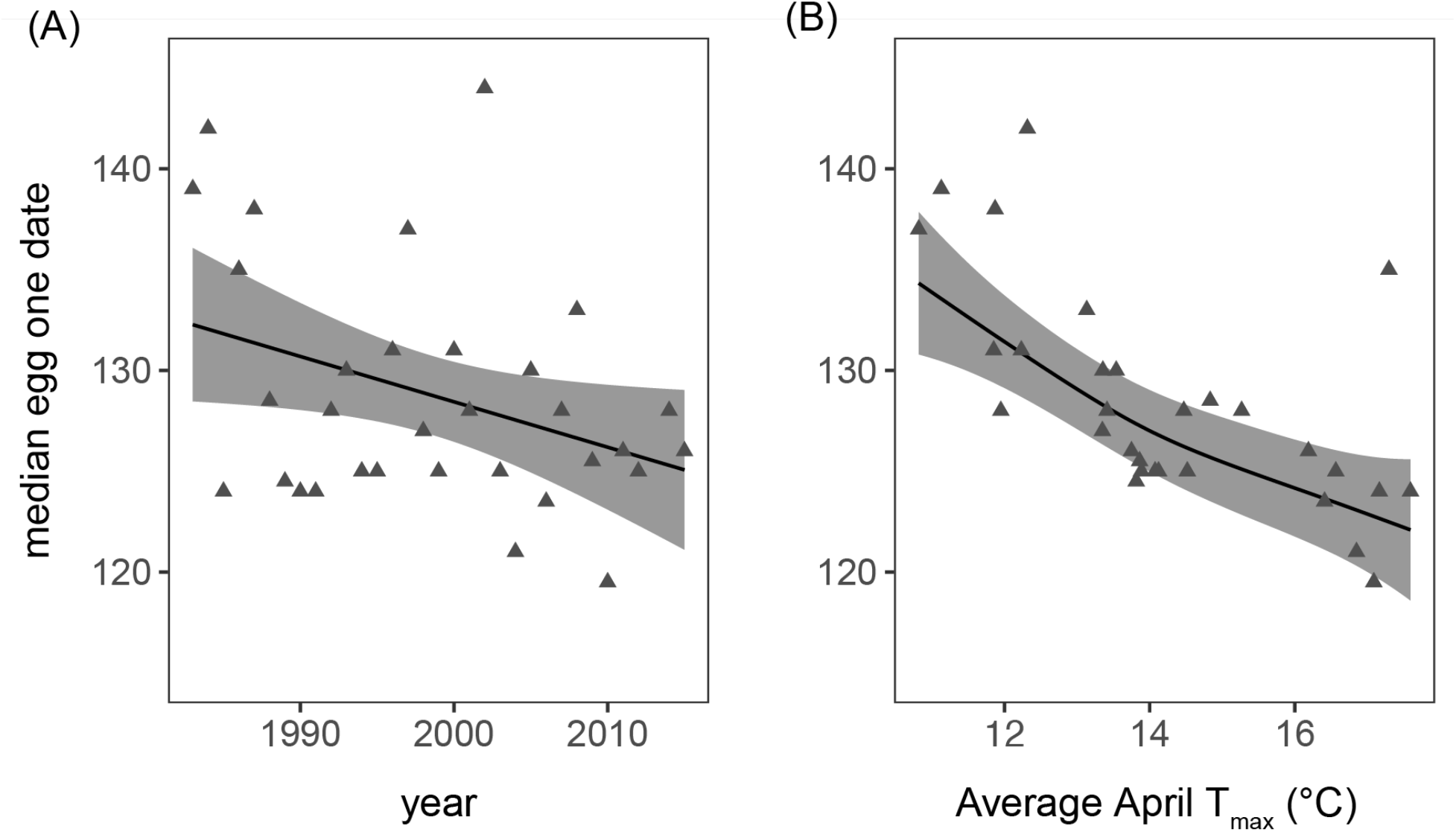
(*A*) Average egg one date of females is shown as a function of year only (*R*^*2*^=0.10). (*B*) Average egg one date is shown as a function of April average maximum temperatures when the model also accounts for nonlinear effects of year (*R*^*2*^= 0.65). Fitted values and 95% confidence bands from the GAMs are overlaid with original data.

### c) Selection analyses

We observed strong selection on egg one date, and the gradient from our full dataset GLMM was significantly negative (i.e., selection favouring earlier breeding; *β*=-0.16, *t*=-4.42, *p*<0.001). This estimate was identical to the selection gradient from an analogous LMM (*β*=-0.16, *t*=-4.91, *p*<0.001). Annual total eggs per female was under positive selection, in which individuals that produced more eggs also had more successful fledglings (*β*=0.05, *t*=5.28, *p*<0.001). Older females had marginally higher relative fitness (*β*=0.12, *t*=1.89, *p*=0.06). Annual nest predation rates did not predict relative fitness (*β*=-0.09, *t*=-0.38, *p*=0.71). The overall (GLMM) selection gradient was similar to the mean of the per-year estimates 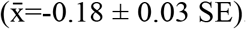. Annual selection gradients showed strong inter-year variation (σ²=0.04) and became significantly more negative (i.e., more strongly favouring earlier breeding) with time (Fig. 3; *β*=-0.01, *p*=0.04, *R*^*2*^=0.11).

**Figure 3.**
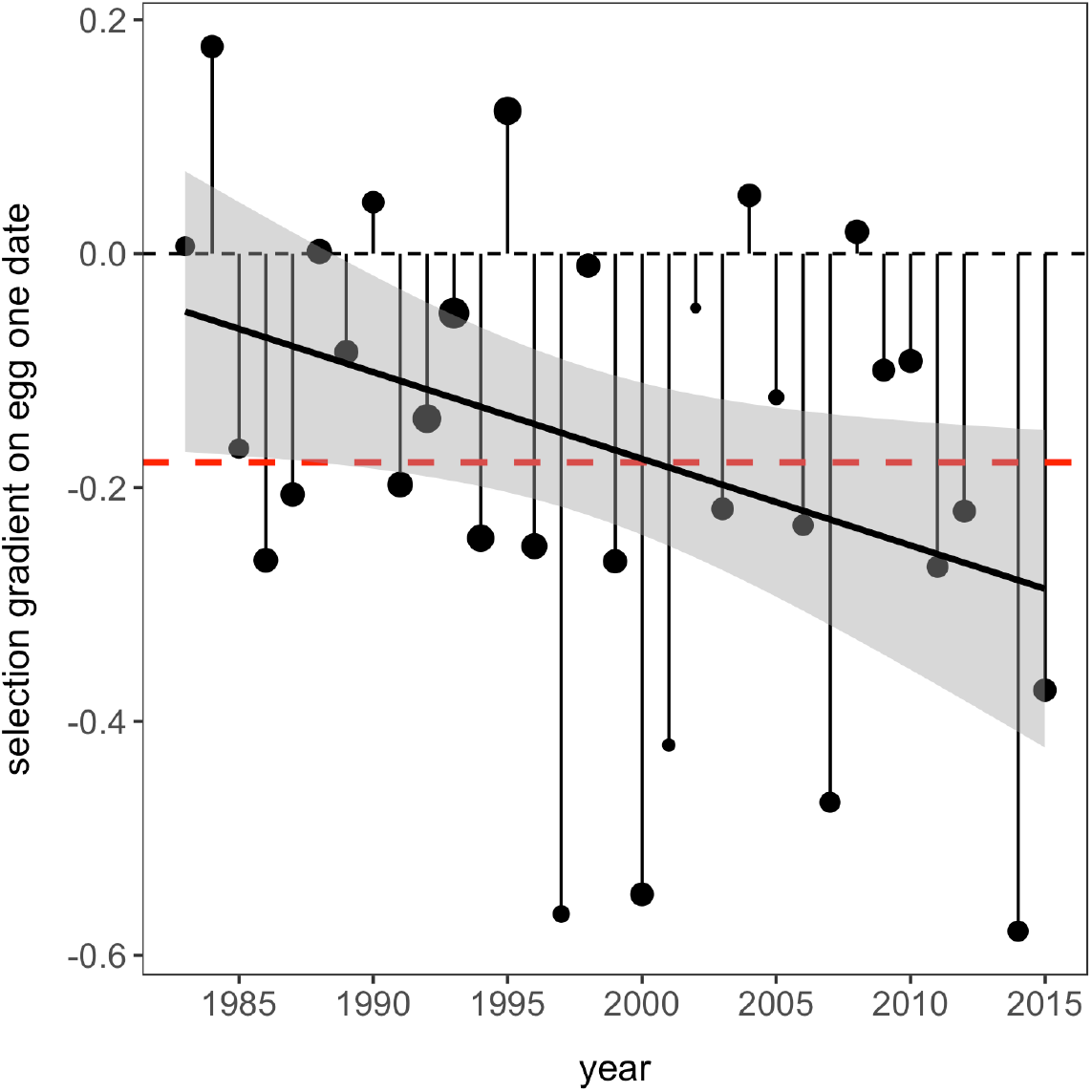
Per-year selection gradients on egg one date estimated with compound Poisson GLMs after adjusting for annual total eggs per female and female age. The dashed line shows β=0, whereas the red line displays the mean selection gradient across the 32 years. The solid line and grey band show fitted values and 95% confidence intervals from a linear model that included weighting by annual sample size (points are scaled by sample size).

When comparing temperature-dependent models of selection on egg one date, all three April temperature measures received equivalent support (Table S3). When analysing selection gradients directly, models including average April T_max_ and T_min_ received the most support from AICc (*w*_*i*_=0.36 and 0.35), but warmer temperatures were nevertheless not associated with selection gradients (T_max_: *β*=0.01, *p*=0.55, *R*^*2*^=0; Fig. 4A; T_min_: *β*=-0.01, *p*=0.59, *R*^*2*^=0). Individual-level GLMMs with interactions between egg one date and temperature did not differentiate between April temperature predictors (*w*_*i*_=0.32–0.35; Table S3). We similarly found no significant interaction between egg one date and April temperature (e.g., T_max_: *β*=-0.0004, *t*=-0.02, *p*=0.98; Fig. 4B). Thus, in both analyses, warmer April temperatures were not associated with selection favouring earlier breeding.

**Figure 4.**
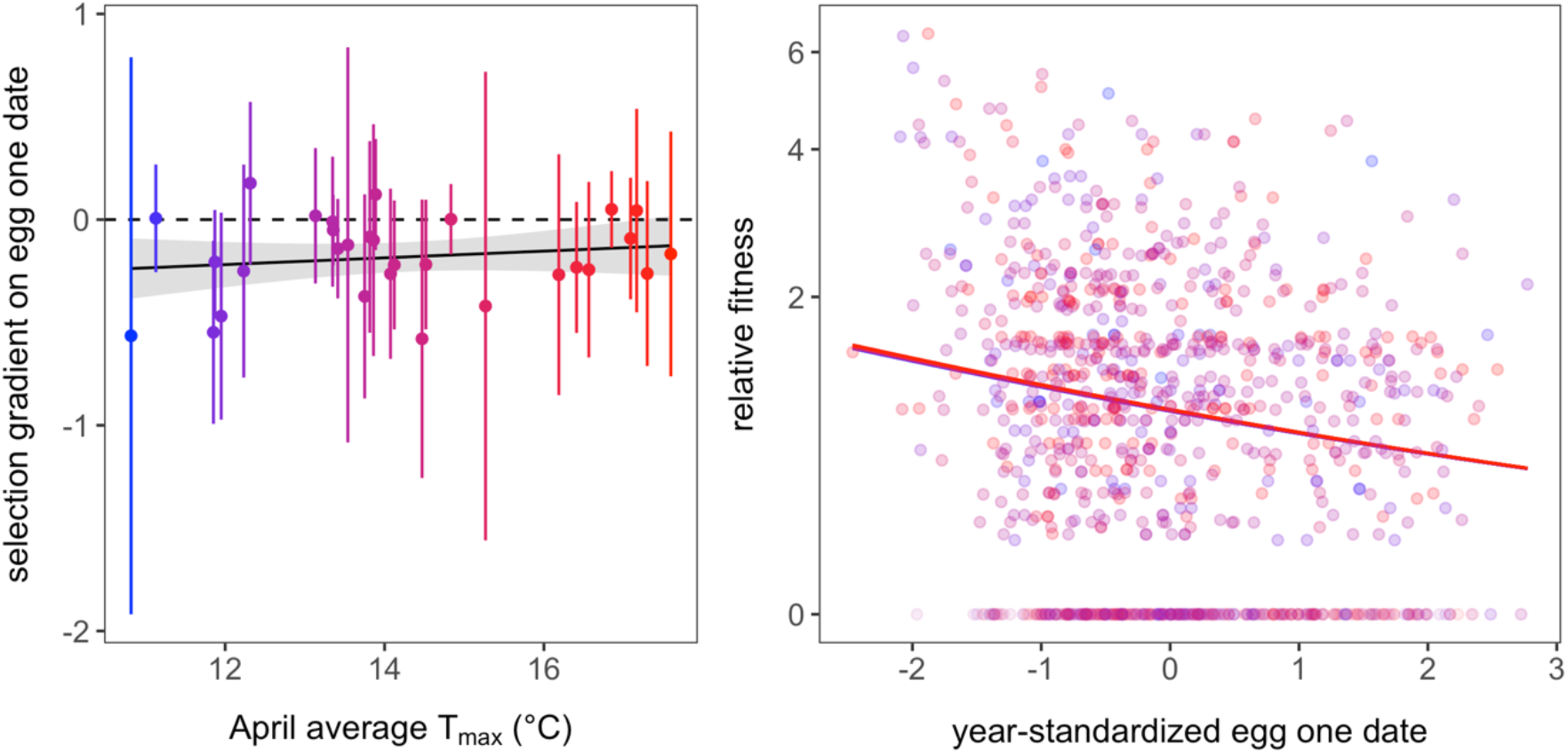
April temperature does not predict variation in selection on egg one date. (A) Results of a linear model using the per-year selection gradients as the response variable (after adjusting for female age and annual total eggs per female), with the solid line and grey band showing fitted values and 95% confidence intervals. The linear model included weighting by annual sample size, which is illustrated through point size. (B) Results of a GLMM predicting relative fitness as a function of the interaction between maximum April temperature and egg one date (after adjusting for female age, annual total eggs per female, and annual predation rate). Points display individual nest data and lines show fitted values, with colours indicating temperature as in A. Relative fitness is shown with a modulus transformation given the right-skew in this variable.

### d) Behavioural plasticity

We used RMMs to assess phenotypic plasticity in the relationship between timing of breeding and temperature. For 62 females studied across at least three years, RRMs found most support for an association between egg one date and April average T_min_ (*w*_*i*_=0.60) and T_med_ (*w*_*i*_=0.36, ΔAICc=1.04), but not with T_max_ (*w*_*i*_=0.04, ΔAICc=5.33; Table S4). Individuals started breeding significantly earlier with warmer April average T_min_ (*β*=-1.33, *t*=-2.50, *p*=0.02) and T_med_ (*β*=-1.67, *t*=-2.70, *p*=0.01). Importantly, individuals did not vary in their elevation (i.e., estimated egg one date at the average temperature) nor their slope (i.e., individual response to inter-year variation in temperature) for either competitive temperature measure (Fig. 5; Table S5). Therefore, females displayed significant population-level phenotypic plasticity, but not inter-individual variation in plasticity.

**Figure 5.**
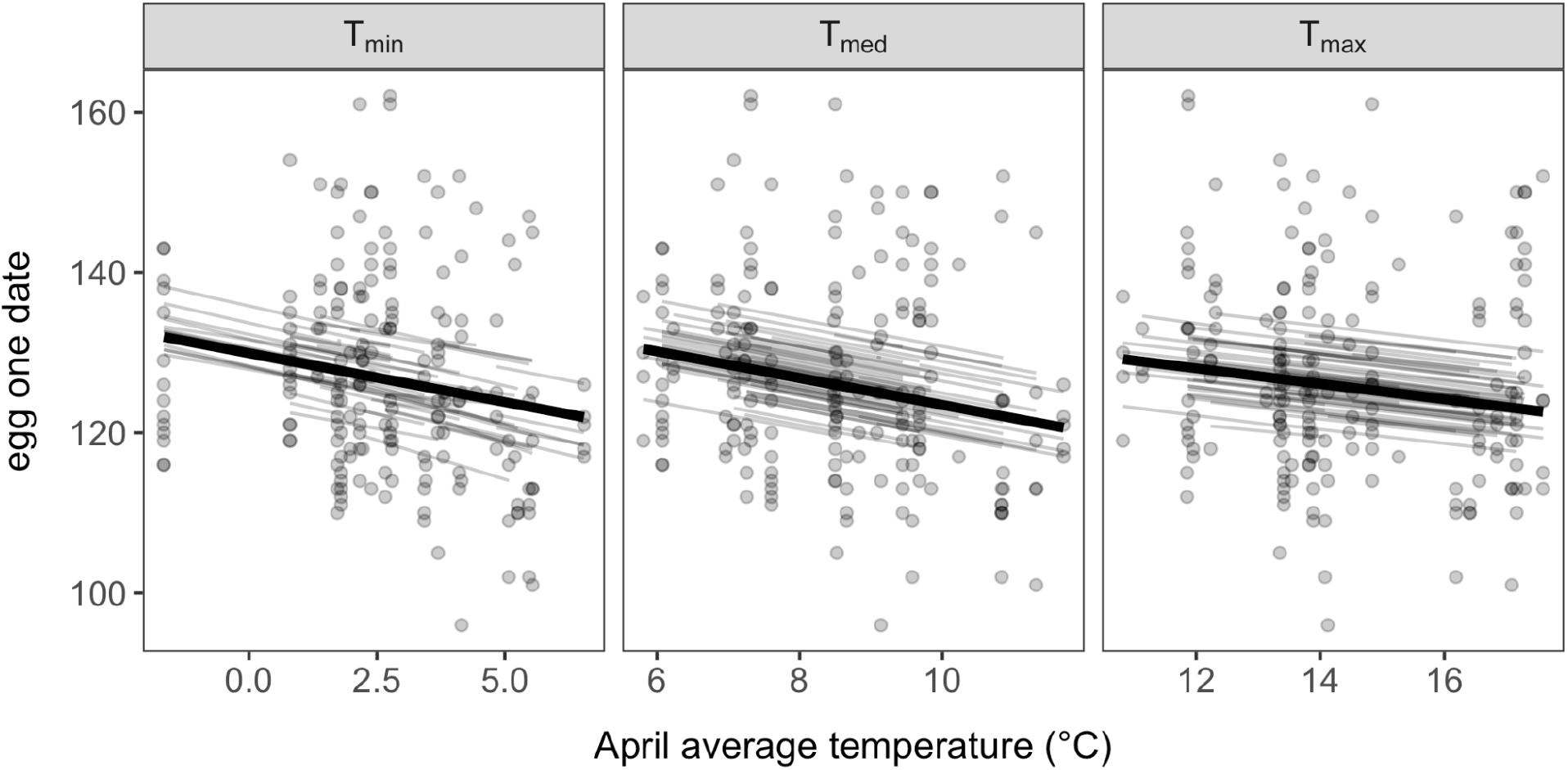
Fitted values of RRMs testing for temperature-driven plasticity in egg one date. Thick lines show the overall reaction norm for each April temperature measure (displayed in order of Akaike weights) after controlling for female age, annual total eggs per female, and annual predation risk, whereas thin lines show reaction norms for each individual female. Random effects were visualized by holding each female (assumed to be ASY) at its mean annual total eggs per female and annual predation rate.

## 4. Discussion

We investigated shifts in phenology over a 32-year period and in relation to spring temperatures and found a net change of 12 days in egg one date and a maximum between-year advance of 24 days. Springs have grown warmer, and females are initiating reproduction earlier than in the past. We also found evidence of selection favouring earlier breeding that has also become stronger over time. However, unlike studies of other avian species [25, 30], spring temperatures did not predict strength of selection on egg one dates, suggesting that other factors may be driving shifts in phenology. Among the many possibilities are some combination of abiotic factors (Dunn & Winkler 2010) or even advances in male reproductive stimulating earlier egg-laying in females (Watts, Edley & Hahn 2016). It is also possible that females respond to a temperature threshold in the spring to initiate laying, explaining why our temperature measures were not related on selection on timing of breeding.

We found strong evidence for plasticity in driving these phenological changes. For a subset of returning females with sufficient multi-year data, females bred earlier in the warmer of the three or more springs in which they bred. However, females exhibited very little inter-individual variation in the degree of plasticity, leaving little variation on which selection might act. While selection and plasticity both likely played a role in earlier breeding associated with warmer springs, the pattern may be more a result of plasticity than selection.

### a) Plastic versus evolutionary responses to climate change

Microevolutionary responses to climate change are predicted to result from directional selection favouring earlier breeding to alleviate the negative effects of phenological mismatches (Charmantier & Gienapp 2014). Without knowledge of the heritability of timing of reproduction, however, evidence of selection favouring earlier breeding is insufficient to conclude that microevolutionary change is occurring, as genetic and environmental effects can be difficult to disentangle (Merilä 2012; Helm *et al*. 2019). There is some evidence of microevolutionary changes in phenology across taxa that are likely adaptive shifts in response to climate change (Van Asch *et al*. 2013; Manhard, Joyce & Gharrett 2017). In the case of the junco, while we found strong overall selection favouring earlier breeding, and this selection has become stronger over time, the strength of selection was not associated with the observed changes in spring temperature. We note that we did not measure genetic variation or heritability of reproductive timing. Therefore, we cannot conclude whether microevolutionary change might account for the observed shifts in timing. Future work should integrate genomic quantitative genetics approaches with this breeding data to determine the role of microevolution in phenological shifts, which could in turn have important implications for conservation genomics (Gienapp *et al*. 2017).

Additionally, behavioural plasticity, which can allow for more rapid changes in phenotype than microevolutionary changes, may account for why earlier breeding was related to increases in fitness over time (Van Buskirk, Candolin & Wong 2012; Charmantier & Gienapp 2014; Beever *et al*. 2017). Numerous studies support behavioural plasticity as a mechanism for coping with climate change (Charmantier *et al*. 2008; Phillimore *et al*. 2016; Verhagen *et al*. 2020), despite its limitations in promoting population persistence in the face of climate change (Ghalambor *et al*. 2007; Gienapp *et al*. 2013; Duputié *et al*. 2015). We found that returning females initiated egg laying earlier in warmer springs. However, there was very little among-individual variation in the degree of plasticity upon which selection might act, suggesting that microevolutionary change in plasticity itself is not a likely explanation for the observed change.

### b) Winners versus losers in relation to climate change

Global change biologists often discuss ‘winners’ and ‘losers’ in relation to climate change, typically in the context of range shifts (Crick 2004; Bateman *et al*. 2016; Tayleur *et al*. 2016). Here, we extend these concepts of winning and losing to migratory strategy and breeding season length. Short-distance migrants and residents often experience longer breeding seasons than long-distance migratory species. This is true in part because they typically breed at lower latitudes where spring comes earlier and also because they do not lose time to the migratory journey leaving time for multiple broods (Newton 2010). Juncos in our study population have a longer breeding season than closely related long-distance migrant populations (Nolan *et al*. 2002). Females can re-nest as many as five times and can fledge up to three successful nests.

The advancement in breeding phenology reported here is supported by a previous finding that multi-brooded species tend to exhibit larger advances in breeding phenology than single-brooded species, likely because multi-brooded species are experiencing longer breeding seasons with warmer springs (Dunn & Møller 2014). Thus multi-brooded populations are expected to have higher reproductive output than migratory populations that are typically single- or double-brooded (Halupka & Halupka 2017), an effect echoed in our finding that juncos that bred earlier tended to fledge more offspring that year, presumably an effect of having more time for breeding attempts (Dunn & Møller 2014). Warmer springs may benefit this population by allowing females to breed earlier and extend their breeding season, despite the lack of evidence that warmer spring temperatures predicted stronger selection favouring earlier breeding.

Overall, females can respond flexibly to changes in temperature, but individuals do not strongly vary in their plastic response to temperatures. However, plasticity alone will likely be insufficient for populations to survive in the long-term when facing climate change (Gienapp *et al*. 2013). Since our study population occurs at high elevation, persistent increases in temperature could eventually result in population decline, as the population cannot shift any further up the mountains.

### c) Future directions

Accurate predictions of future responses to climate change will require further consideration of mechanisms of female reproductive timing (Williams 2012; Chmura, Wingfield & Hahn 2020; Kimmitt 2020). Past and ongoing work in the junco is elucidating the physiological mechanisms driving reproductive timing in females based on life history, including endocrine systems and costs of early breeding (Greives *et al*. 2016; Graham *et al*. 2019; Kimmitt *et al*. 2019; Kimmitt, Sinkiewicz & Ketterson 2020). However, more research is necessary to understand how females integrate supplementary cues, such as temperature, to regulate the final stages of their reproductive development and ovulation (Wingfield *et al*. 2016; Chmura, Wingfield & Hahn 2020). Via our analysis of this 32-year dataset, we found that flexibility in female timing is likely relevant for population persistence, and further work on the proximate mechanisms of female timing will improve forecasts on the effects of climate change on birds.

## Acknowledgments

We acknowledge the significant role of the late Val Nolan Jr., who established this field site for long-term study as a co-PI with Ellen Ketterson in the early 1980s. We also thank Ketterson lab members and colleagues who have contributed to this long-term dataset, with a special thanks to long-term lab managers and field assistants, Eric Snajdr, Charles Ziegenfus, and Sarah Wanamaker. We also acknowledge support from MLBS and its directors, Henry Wilbur and Edmund “Butch” Brodie III, as well as MLBS staff. Finally, we thank Allie Byrd, Alex Jahn, Katie Talbott, and Sarah Wanamaker for discussion and feedback on the manuscript.

## Authors’ Contributions

AAK conceived and designed the study, curated data, conducted statistical analysis and was the primary author of the manuscript; DJB conducted statistical analysis, drafted sections of the manuscript, and revised the manuscript; SND participated in data curation and analysis and revised the manuscript. NMG conceived and designed the study, curated data, and revised the manuscript. KAR conceived and designed the study and revised the manuscript. EDK conceived and designed the study, secured funding for data collection, and revised the manuscript. All authors gave final approval of this publication.

## Data Accessibility

Data will be made available on Dryad pending manuscript acceptance.

