## Supplementary Materials for "Plasticity in female timing may explain current shifts in breeding phenology of a North American songbird"

*Annual search effort*

Researchers typically began monitoring the population in mid-April, with systematic nest searching around May 15 regardless of year; however, research effort varied annually. Therefore, we investigated if changes in breeding phenology were an artifact of search effort. We examined whether egg one dates were normally distributed per year, as a left-skew could indicate bias toward late egg one dates and that our team inadvertently missed the beginning of the breeding season. We tested for annual normality of egg one dates using Shapiro-Wilk tests in R. Egg one data were normal in 19 of 32 years (59.4%), and non-normal years were neither left-skewed nor time-biased (Fig. S1). Additionally, when nests were found at the egg or nestling stage, egg-one date was back calculated based on hatch date or nestling size.

*Comparison of temperature loggers*

All data for the temperature loggers can be accessed via https://mlbs.virginia.edu/meteorological-data.

We used linear models to confirm that temperature data collected by both Logger A and Logger B between March–May in the years 1994–1997 were correlated before combining the datasets. Maximum temperature data from Logger A were left-skewed, however, median and minimum data from both datasets as well as maximum temperature data from Logger B were normally distributed. There was no transformation that would correct the normality of maximum temperature from Logger A. However, using the Breusch Pagan test for Heteroskedasticity from the R package *'olsrr'*, we found that the variance was constant for all three models (minimum temperature: $X^2$=0.06, $p$=0.801; median temperature: $X^2$=2.04, $p$=0.153; maximum temperature:

$X^2$=0.03, $p$=0.865). We found that all temperatures were highly correlated between Logger A and Logger B (Fig. S2; *minimum temperature:* $R^2$= 0.56, $p$< 0.000; *median temperature:* $R^2$= 0.58, $p$< 0.0001; *maximum temperature:* $R^2$= 0.42, $p$< 0.0001).

Thus, for our formal analysis, we combined the datasets: data from Logger A was used from 1983–1994 and data from Logger B was used from 1995–2015.

*Female age as a predictor of relative fitness*

Female age is a known predictor of female lay date in dark-eyed juncos (Bauer et al. 2018). We analyzed the relationship between female age and egg one date using a linear mixed model fit with REML. Females were grouped into two age categories of young females in their first breeding season and old females in a returning (second or later) breeding season. We included both year and female ID as random effects to control for pseudoreplication. Female age predicted egg one date, as old females (i.e., returning females in her second or later breeding season) laid on average 2.7 days earlier than young females (i.e., females in their first known breeding season and classified as first-year females based on plumage coloration) when controlling for both year and female ID (Fig. S3; $t$= -3.97, $p$< 0.0001).

***References***

Bauer, C. M., Graham, J. L., Abolins-Abols, M., Heidinger, B. J., Ketterson, E. D., & Greives, T. J. (2018). Chronological and biological age predict seasonal reproductive timing: an investigation of clutch initiation and telomeres in birds of known age. *The American Naturalist*, *191*(6), 777-782.

Perrins, C. M., & McCleery, R. H. (1989). Laying dates and clutch size in the great tit. The

Wilson Bulletin, 101, 236–253.

Wood, S. (2017) Package “mgcv”: mixed GAM computation vehicle with GCV/AIC/REML smoothness estimation. Version.

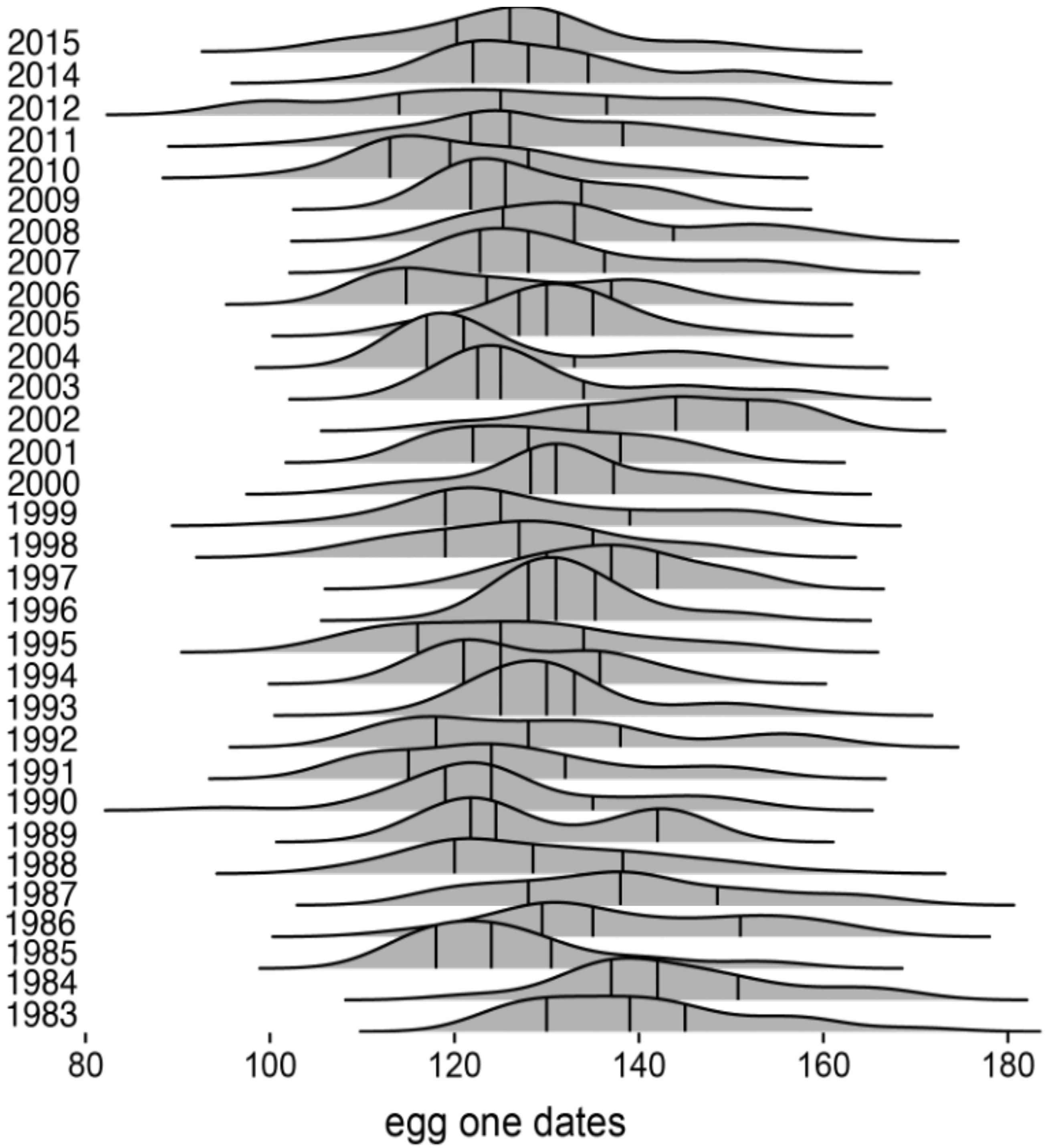


Figure S1. Density distributions of egg one dates for each year. Lines on each density distribution mark the first, second and third quartile.

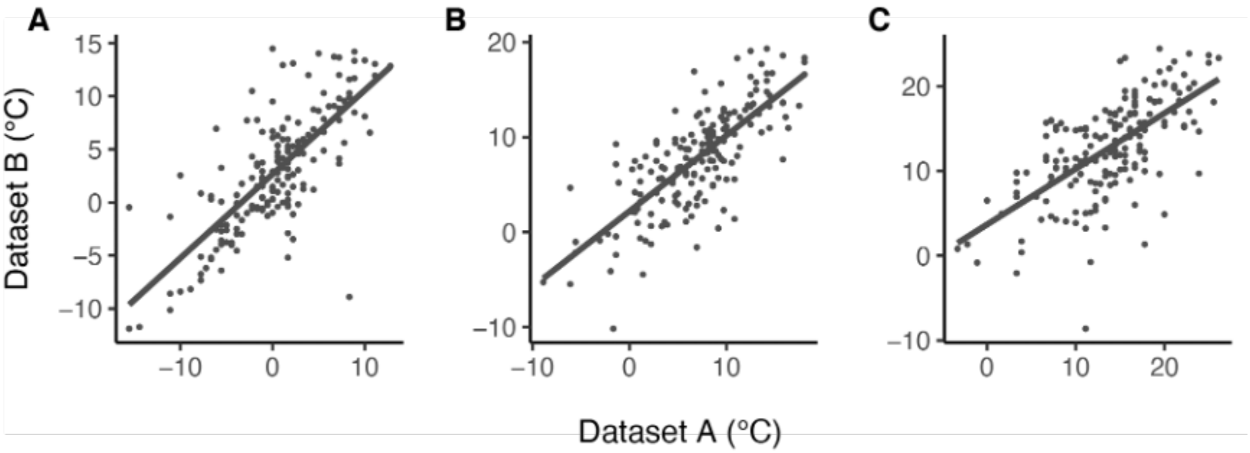


Figure S2. Correlation between temperate data collected from the NOAA weather station (Logger A) and the weather station (Logger B) in March through May in 1994-1997 for (A) minimum temperatures, (B) median temperatures calculated from minimum and maximum temperatures, and (C) maximum temperatures.

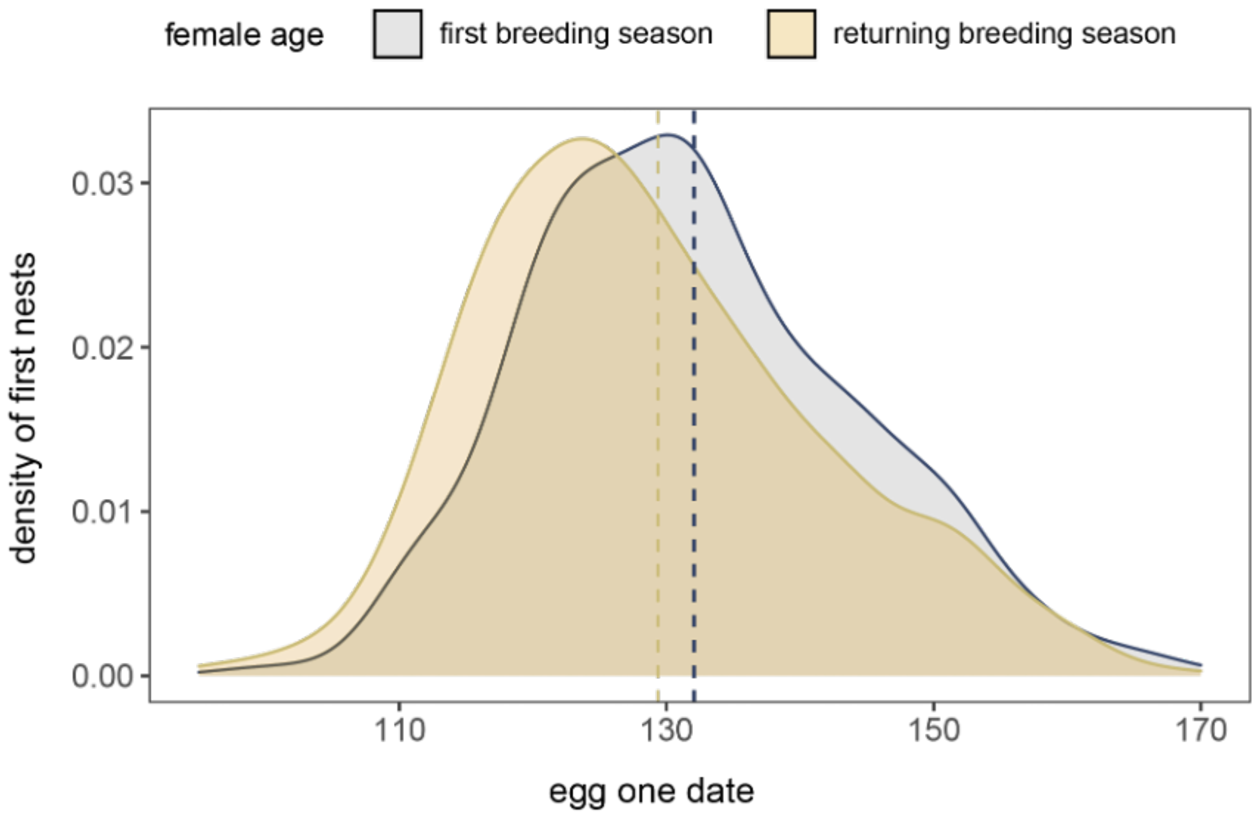


Figure S3. Distribution density of egg one dates for first nests of females by age: females in their first breeding season (i.e., second years) versus females in a returning breeding season (i.e., after second years). Dotted lines are the predicted mean egg one dates for each age group from the GLMM after controlling for year and female ID. Old females lay 2.7 days earlier than young females when controlling for year and female ID.

Table S1. Summary of GAMs testing relationship between year and average monthly median, minimum, and or maximum temperatures.

| Month | Average Monthly $T_{med}$ | | | Average Monthly $T_{min}$ | | | Average Monthly $T_{max}$ | | |
|---|---|---|---|---|---|---|---|---|---|
| | $F_{1,1}$* | *P-value* | $R^2$ | $F_{1,1}$ | *P-value* | $R^2$ | $F_{1,1}$* | *P-value* | $R^2$ |
| March | 0.004 | 0.95 | 0.00 | 1.47 | 0.236 | 0.02 | 0.05 | 0.833 | 0.00 |
| April | **4.79** | **0.037** | **0.11** | **15.86** | **<0.001** | **0.33** | 0.18 | 0.679 | 0.00 |
| May | 1.56 | 0.158 | 0.08 | **11.41** | **0.002** | **0.25** | 1.45 | 0.292 | 0.07 |

*May $T_{med}$: $F_{1.41, 1.71}$; May $T_{max}$: $F_{2.08, 2.60}$

Table S2. Comparison of GAMs testing relationships between median egg one dates and temperatures while accounting for nonlinear effects of year. Models are ranked by ΔAICc alongside the Akaike weights ($w_i$) and adjusted $R^2$.

| Fixed effects | ΔAICc | $w_i$ | $R^2$ |
|---|---|---|---|
| s(Average April $T_{max}$) + s(year) | 0.00 | 0.97 | 0.63 |
| s(Average April $T_{med}$) + s(year) | 7.32 | 0.03 | 0.55 |
| s(Average April $T_{min}$) + s(year) | 17.93 | 0.00 | 0.53 |
| s(Average March $T_{max}$) + s(year) | 18.83 | 0.00 | 0.19 |
| s(Average May $T_{min}$) + s(year) | 19.26 | 0.00 | 0.18 |
| s(Average March $T_{med}$) + s(year) | 19.50 | 0.00 | 0.17 |
| s(Average May $T_{med}$) + s(year) | 19.71 | 0.00 | 0.16 |
| s(Average March $T_{min}$) + s(year) | 19.99 | 0.00 | 0.16 |
| s(Average May $T_{max}$) + s(year) | 20.00 | 0.00 | 0.16 |

Table S3. Comparison of linear models predicting selection gradients on egg one date (A) and GLMMs predicting relative fitness by the interaction of egg one date and each temperature covariate (B). Models are ranked by ΔAICc alongside the Akaike weights ($w_i$) and $R^2$ (adjusted $R^2$ for linear models, $R^2_m$ and $R^2_c$ for GLMMs). Linear models include weighting by sample size, and GLMMs include random effects of year and ID.

| (A) Linear models of selection gradients | ΔAICc | $w_i$ | $R^2_m$ | $R^2_C$ |
|---|---|---|---|---|
| Average April $T_{max}$ | 0 | 0.36 | 0.0 | NA |
| Average April $T_{min}$ | 0.07 | 0.35 | 0.00 | NA |
| Average April $T_{med}$ | 0.38 | 0.30 | 0.00 | NA |
| (B) GLMMs of full dataset (fixed effects only) | ΔAICc | $w_i$ | $R^2_m$ | $R^2_C$ |
| Egg one date * average April $T_{min}$ + clutch + age + depredation | 0 | 0.35 | 0.07 | 0.13 |
| Egg one date * average April $T_{med}$ + clutch + age + depredation | 0.09 | 0.33 | 0.07 | 0.13 |
| Egg one date * average April $T_{max}$ + clutch + age + depredation | 0.16 | 0.32 | 0.07 | 0.13 |

Table S4. Comparison of RRMs testing for temperature-driven plasticity in egg one date within and between individual females. Models are ranked by ΔAICc alongside the Akaike weights ($w_i$) and both $R^2_m$ and $R^2_c$.

| RRM structure | ΔAICc | $w_i$ | $R^2_m$ | $R^2_C$ |
|---|---|---|---|---|
| Average April $T_{min}$ + clutch + age + predation + (1\|year) + (Average April $T_{min}$\|female ID) | 0 | 0.60 | 0.20 | 0.29 |
| Average April $T_{med}$ + clutch + age + predation + (1\|year) + (Average April $T_{med}$\|female ID) | 1.04 | 0.36 | 0.21 | 0.26 |
| Average April $T_{max}$ + clutch + age + predation + (1\|year) + (Average April $T_{max}$\|female ID) | 5.33 | 0.04 | 0.19 | 0.27 |

Table S5. Estimates of variance for all random effects in the most competitive RRMs (fitted with REML) assessing temperature-driven plasticity in egg one date. Significant differences from zero were tested using sequential likelihood ratio tests.

| | $T_{min}$ | | | $T_{med}$ | | |
|---|---|---|---|---|---|---|
| Term | $\sigma^2$ | LRT | *p* | $\sigma^2$ | LRT | *p* |
| Female ID | 0.00 | 0.10 | 0.75 | 0.04 | 0.09 | 0.76 |
| Average April temperature\|female ID | 0.63 | 1.07 | 0.58 | 0.004 | 0.14 | 0.93 |
| Year | 6.44 | 0.70 | 0.40 | 7.40 | 0.73 | 0.39 |
| Residual | 110.56 | | | 116.41 | | |
